# Regulon-informed cellular representations reveal task-dependent generalization in drug combination prediction

**DOI:** 10.64898/2026.09.23.753774

**Authors:** Elizaveta Ignatova, Maksim Likhter

## Abstract

Drug combination models must represent cellular context in a form that remains informative when the tested cells or compounds differ from those used for training. Transcription factor regulons offer a biologically structured representation, but their contribution can depend on the accompanying features and the intended prediction task. We developed a symmetric three-class predictor of DrugComb ZIP interactions and compared all seven combinations of landmark expression, Hallmark pathway scores and CollecTRI regulon activities. The common dataset contained 302,042 observations across 3,042 drugs and 155 cellular contexts. We evaluated three group-held-out generalization settings: unseen cell lines, unseen drug scaffolds, and simultaneous exclusion of both, with five training seeds per representation and partition. Pathways plus regulons achieved the highest mean Macro-F1 on unseen cell lines (0.4737, compared with 0.4503 for expression alone). Compared with pathways alone, adding regulons increased mean Macro-F1 by 0.0153 on unseen cell lines and 0.0229 on unseen drug scaffolds, with positive differences across all five paired seeds in both settings. In contrast, the same addition reduced the mean under joint context and scaffold exclusion. Representation rankings also depended on the objective: regulons alone achieved the highest synergy average precision on unseen lines, whereas the Macro-F1-leading combination did not maximize precision among the top-ranked candidates. Further compression of pathways into non-negative matrix factorization programs did not exceed the best context-model Macro-F1 in any setting. These results demonstrate that the predictive value of functional context representations depends on both the type of domain shift and the evaluation objective, and that combining additional biological representations does not necessarily improve generalization. They provide a computational basis for studying drug combinations in senescent states, where transfer must subsequently be assessed using state-specific responses and matched controls.

## Introduction

The activity of a drug combination depends on both the compounds and the biological system in which they are tested. Large combination screens make it possible to learn associations between chemical and cellular features and experimental interaction scores. Resources such as DrugComb support this work by harmonizing combination measurements from multiple studies (Zheng et al. 2021). However, the utility of a predictive model depends on its intended setting. Recovering measurements involving familiar cells and compounds, prioritizing new compounds in familiar cells, and predicting responses in an unfamiliar cellular context are distinct tasks. A representation that performs well in one setting need not retain its advantage in another.

Neural approaches have explored chemical fingerprints, molecular graphs, target information and gene expression, with different strategies for combining these inputs. DeepSynergy, MatchMaker and TranSynergy exemplify developments in feature integration and context-dependent prediction (Preuer et al. 2018; Kuru et al. 2022; Liu and Xie 2021). Evaluations of this field have also shown the importance of preprocessing, feature choices, simple baselines and data-splitting procedures (Baptista et al. 2023; Tasnina et al. 2025). These observations make controlled comparisons of cellular representations valuable even when the downstream predictor is held largely constant. Such comparisons ask which information supports a particular form of generalization, rather than attributing every performance change to a new architecture.

Regulon activities are one candidate representation. A regulon links a transcription factor to its target genes, allowing a coordinated transcriptional pattern to be summarized as an activity score. CollecTRI supplies signed transcription factor–target interactions, and decoupleR provides methods for estimating activities from molecular measurements (Müller-Dott et al. 2023; Badia-i-Mompel et al. 2022). Pathway gene sets provide a complementary description at the level of coordinated biological processes (Liberzon et al. 2015). Both representations encode prior biological organization, but they summarize different and partly overlapping aspects of the same expression profile. Whether these representations improve interaction prediction individually or in combination, and whether their relative value changes under different forms of domain shift, are empirical questions.

The use of transcription factor information in this task has a direct precedent. CCSynergy compares several cellular representations, including a transcription factor representation obtained from CARNIVAL-inferred signaling networks, and evaluates generalization to held-out tissues and combinations (Hosseini and Zhou 2023). Its Supplementary Methods S2 describes a categorical representation of 100 transcription factor nodes. Here, we investigate a different operational representation: continuous activities for 771 transcription factors inferred from each basal expression profile using the same fixed CollecTRI regulatory prior. We systematically evaluate these activities alone and in all combinations with landmark expression and Hallmark pathway scores. This design makes it possible to distinguish the contribution of adding regulons to a given feature set from the performance of an unrelated model using different inputs.

A longer-term motivation is prediction in cellular states associated with aging. Senolytic research illustrates why the cellular state matters: the desired effect is selective elimination of senescent cells while preserving appropriate controls (Zhu et al. 2015). Senescence itself is heterogeneous and requires multiple complementary markers for its characterization (Gorgoulis et al. 2019). SenolyticSynergy has already linked oncology combination learning to this application through aging-associated genomic features and candidate senolytic combinations (Ye et al. 2025). The central challenge for a transferable representation is to retain information relevant to response when the underlying cell state changes. A functional representation provides a way to formulate this challenge, while direct response measurements are needed to determine whether the transfer succeeds.

Here, we present a context-conditioned predictor and systematically compare seven cellular-context representations across three generalization settings: held-out cell lines, held-out drug scaffolds, and simultaneous exclusion of both. The primary factorial experiment comprises 105 training runs evaluating all combinations of landmark expression, Hallmark pathway activity, and CollecTRI-derived transcription-factor activity across five training seeds. In a separate analysis, we further examine whether compressing pathway activities into low-dimensional NMF-derived programs alters generalization, using 300 candidate training runs with program dimensionality selected on validation data. We evaluate overall three-class classification performance, class-specific prediction, candidate-ranking performance, and variability across held-out cellular contexts. Together, the analyses reveal task-dependent complementarity among biological representations: the representation that performs best depends on the type of domain shift and evaluation objective, while combining additional features or increasing the level of abstraction does not uniformly improve generalization.

## Methods

### Dataset assembly and interaction labels

We used a DrugComb summary snapshot containing 739,964 rows. Measurements were aggregated by unordered drug pair, cell line and study, with replicate measurements summarized within, but not across, studies, producing 639,410 observations. The target was the median zero-interaction potency (ZIP) score within each aggregate. ZIP measures departure from a reference model of non-interaction (Yadav et al.

2015). We assigned antagonism to values below −10, synergy to values above +10, and no interaction to the closed interval [−10, +10]. The latter label therefore denotes a score interval and does not imply absence of single-agent activity.

Drug identifiers were mapped to chemical structures and target features, and cellular identifiers were mapped to DepMap models. An observation was eligible only when both compounds had chemical features and entries in the target, propagated-network and target-availability matrices, and the cellular context had available landmark-expression, Hallmark-pathway, and CollecTRI-derived transcription-factor activity features. A drug could have a valid all-zero target or propagated vector when no known targets were available or when known targets did not map to the interaction network; a three-state availability indicator distinguished these cases from drugs with network-mapped targets. The same eligibility filter was applied to every context representation, ensuring that all models were evaluated on the same observations.

The resulting dataset contained 302,042 observations, 301,069 distinct pair–context identities, 3,042 drugs and 155 cellular contexts. Classes comprised 25,624 antagonistic, 242,193 no-interaction and 34,225 synergistic observations. Study-specific aggregates of the same pair and context remained separate observations. Among 973 pair–context identities with multiple eligible observations, 328 had discordant interaction labels across studies. Identical pair–context identities were kept within a single partition. Neither study identifiers nor interaction labels were used as input features. Doses and exposure durations were not explicit inputs to the classifier.

### Drug features and shared encoding

Chemical structures were represented using radius-2 (ECFP4) Morgan fingerprints with 2,048 bits and chirality enabled (Rogers and Hahn 2010). The broader structure resource contained 8,192 valid SMILES among 8,396 DrugComb identifiers. We used a fixed target dictionary and mapped target genes to a HuRI-derived network of 8,245 proteins and 52,068 undirected edges (Luck et al. 2020). Each drug received a binary direct-target vector and a continuous propagated profile in the same protein order.

Target propagation used random walk with restart (RWR). With row-normalized adjacency matrix W, row seed vector s, and restart probability r = 0.5, the propagated profile was updated as *p*^(*t* +1)^ =(1 *−r*) *p*^(*t*)^*W* + *r s* until convergence at a tolerance of 10^−6^. Seed mass was distributed equally among mapped targets. Three mutually exclusive availability indicators distinguished drugs with no known targets, known targets outside HuRI, and at least one target in HuRI. In the full dictionary, 4,218 drugs had known seeds and 2,343 had at least one HuRI-mapped target. These coverage counts describe the resource, whereas analyses of eligible observations used the actual two-drug coverage of each pair.

The same drug encoder, with shared weights, was applied to both compounds. The chemical branch used dimensions 2,048→512→128, the direct-target branch 8,245→256→32, and the propagated branch 8,245→512→128. Their outputs were concatenated with the three availability indicators to form a 291- dimensional representation, which was passed through a 291→256→128 fusion network to produce a 128- dimensional intrinsic drug embedding. The drug encoder was trained jointly with the interaction-prediction objective. All context comparisons used the same drug-feature definitions and drug-encoder architecture.

### Cellular representations

We used basal DepMap protein-coding expression in log²(TPM+1) units and selected the default profile for each model. DepMap and CCLE characterize cellular models across multiple molecular modalities (Ghandi et al. 2019). The expression snapshot contained 1,719 default profiles. A direct audit of its 19,215 gene columns found no nonfinite expression values in these profiles. The preprocessing code supports gene-median imputation, but no such replacements were required for the audited input.

Three feature sets were constructed. **Expression (E)** comprised 942 measured genes from the external set of 978 L1000 landmark genes (Subramanian et al. 2017). **Pathways (P)** comprised univariate linear model (ULM) scores for 50 MSigDB Hallmark gene sets. All retained membership weights were +1; the measured network contained 7,295 pathway–gene memberships. **Regulons (R)** comprised ULM activities for a fixed CollecTRI regulatory network containing 771 factors and 41,714 retained signed TF–target interactions. Gene sets and regulons were required to contain at least five measured targets.

ULM fits an expression profile against the weights of one gene set or regulon across genes within each sample and returns the t-statistic of the fitted slope (Badia-i-Mompel et al. 2022). Thus, each cellular context receives its own continuous activity scores under a common fixed prior. Hallmark scoring used its measured gene-set universe, whereas CollecTRI scoring used the retained signed TF–target interactions represented in the DepMap expression matrix. The regulon expression universe contained 6,836 genes. Because the DepMap values were already provided as log²(TPM+1), no additional logarithmic transformation was applied. Gene orders and the exact retained priors were saved with the experiment.

The seven representations were E, P, R, E+P, E+R, P+R and E+P+R. Each active representation was encoded into a 64-dimensional branch: E used dimensions 942→256→64, P used 50→64→64, and R used 771→256→64. Active branch outputs were concatenated, yielding 64, 128, or 192 features in models containing one, two, or three representations, respectively, and passed through a 256-unit fusion layer to produce a 128-dimensional cellular-context embedding. Within each split, input features were standardized using means and standard deviations estimated across unique training contexts; the resulting parameters were then applied unchanged to validation and test contexts. The fixed CollecTRI regulatory network used for the 105-run comparison did not incorporate correlations estimated across DepMap models.

### Context conditioning and interaction prediction

The 128-dimensional cellular-context vector conditioned each 128-dimensional intrinsic drug embedding using feature-wise linear modulation (FiLM), an explicit drug–context interaction module and a gated residual connection. The same context-conditioning module, with shared weights, was applied to both drugs. The resulting conditioned embeddings of drugs A and B were then combined using their elementwise sum, absolute difference and elementwise product, and concatenated with the context vector. These four 128- dimensional components produced a 512-dimensional pair representation, which was passed through a 512→256→128→3 classifier. The shared drug encoder and context-conditioning module, together with the symmetric pair operations, made the resulting prediction invariant to the ordering of drugs A and B. Softmax outputs provided scores for the three interaction classes, and the class with the highest probability was used as the predicted label.

Networks used GELU nonlinearities, LayerNorm and dropout of 0.20. The number of trainable parameters ranged from 8,248,035 for P to 8,752,995 for E+P+R. The P+R model contained 8,478,627 parameters, close to the 8,498,467 parameters of E. Supplementary Tables S1 and S2 specify branch dimensions and parameter counts.

**Figure 1.**
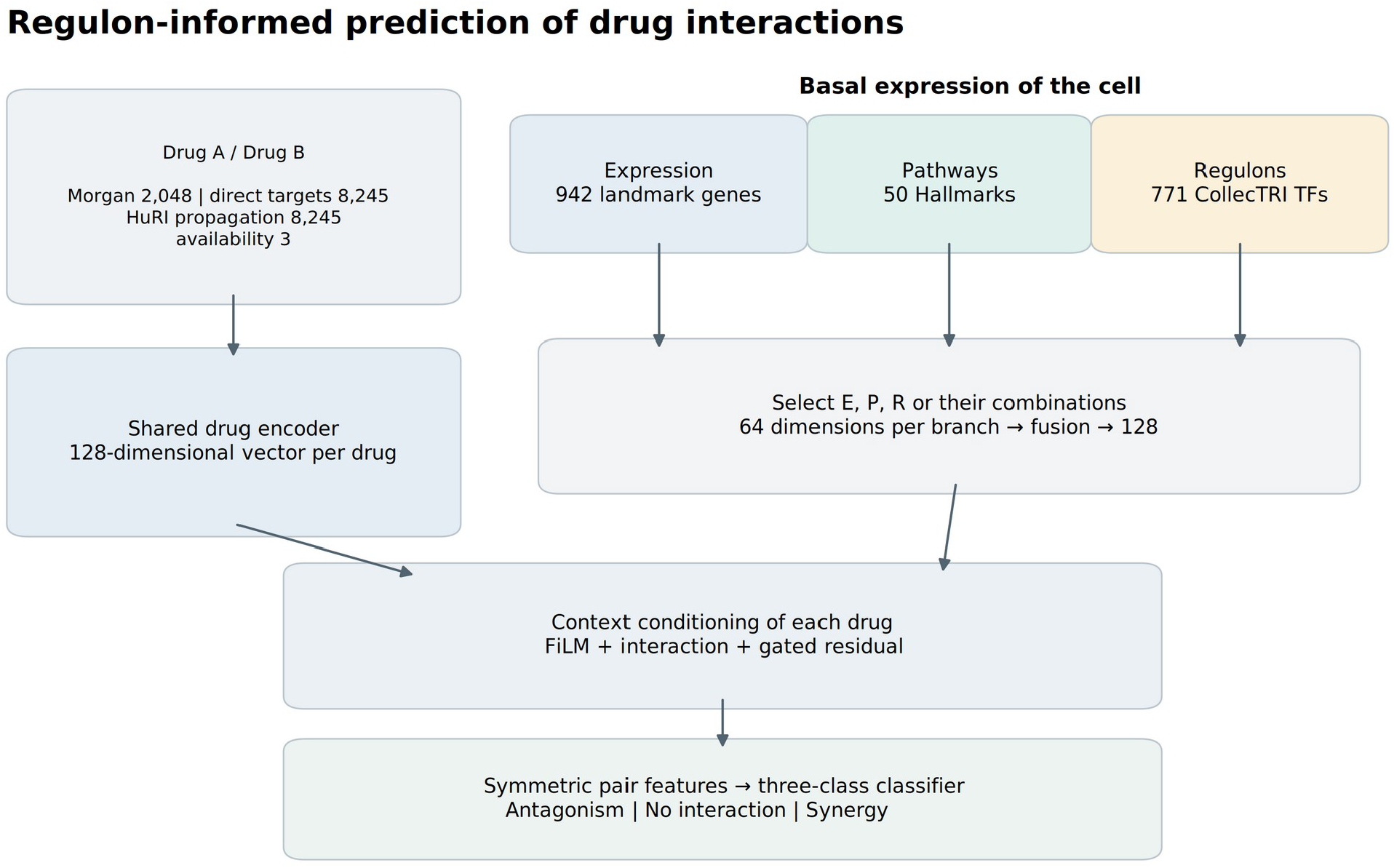
Model architecture. Both drugs are processed by a shared drug encoder. E, P and R denote landmark expression, Hallmark pathway activity and fixed CollecTRI-derived transcription-factor activity, respectively. Each selected cellular-context branch produces a 64-dimensional representation, and active branches are fused into a 128-dimensional context vector. The context vector conditions each drug embedding before symmetric pair construction using the sum, absolute difference and elementwise product of the conditioned drug embeddings. The resulting representation predicts three interaction classes—antagonism, no interaction and synergy—defined by ZIP thresholds described in Methods. The diagram shows the architecture used throughout the cellular-context representation experiments.

### Group-held-out evaluation

We evaluated three forms of held-out information. In **unseen cell line**, complete DepMap model identities were assigned to training, validation or test partitions. In **unseen drug**, compounds were grouped by RDKit Bemis–Murcko scaffolds computed without chirality. Acyclic compounds yielding empty scaffolds were grouped by their full canonical structures. Training contained only pairs in which neither drug belonged to a validation or test drug group. Validation observations required at least one validation-group drug and no test-group drug, whereas test observations required at least one test-group drug and no validation-group drug. Thus, a held-out drug observation required at least one structurally held-out compound, rather than requiring both compounds to be unseen. In **unseen both**, those drug-group rules were combined with independently assigned cellular context partitions, such that test observations required both a test context and at least one test-group drug. Cross-partition combinations were discarded.

Consequently, a drug-held-out test pair could contain a familiar partner. The joint setting required both a held-out cellular context and at least one held-out drug group. We reconstructed the assignments from the retained chemical structures and split metadata, verified the recorded partition sizes and test membership, and confirmed that canonical structures assigned to held-out drug groups did not occur among training drugs. Pair–context identities did not cross partitions.

The split-search procedure considered 50 candidate assignments and selected partitions according to target partition sizes and class proportions, requiring all three interaction classes to be represented in every partition. Interaction labels were used to balance class distributions across partitions but not to select splits according to predictive performance. For training seeds 42–46, the procedure selected identical data assignments within each evaluation setting, corresponding to split seeds 79, 47 and 60 for cell-line, drug and joint holdouts, respectively. The five runs per representation therefore quantify variation arising from model training on one fixed data partition within each generalization setting.

### Training and analysis

Each representation was trained with five seeds in each setting, yielding 105 runs. Optimization used AdamW with learning rate 10^−4^, weight decay 10^−4^ and batch size 128. Weighted cross-entropy used class weights N/(3n_c) estimated from the training partition. Training lasted at most 50 epochs, with early stopping after eight epochs without improvement in validation Macro-F1. ReduceLROnPlateau used factor 0.5 and patience 2, and gradient norms were clipped at 5. The checkpoint with the highest validation Macro-F1 was evaluated on test. Mixed precision was enabled on CUDA, with deterministic CUDA algorithms not enforced.

Macro-F1 was the primary metric, calculated from argmax predictions over the three classes. Additional metrics were balanced accuracy, macro one-versus-rest AUROC, and per-class average precision (AP). The field named AUPRC in the original output was implemented using average_precision_score; we therefore use AP when describing it. Means and sample standard deviations summarize the five runs. We independently recomputed the metrics from saved test probabilities and checked all 105 files against their identifiers, labels and run summaries.

To isolate the operational contribution of adding regulons, we evaluated E→E+R, P→P+R and E+P→E+P+R at matched training seeds. Exact two-sided Wilcoxon tests were applied to the five paired Macro-F1 differences. Holm correction covered the nine contrasts formed by three additions and three settings. These tests concern training-seed variation under the retained assignments; they do not estimate variation across alternative held-out partitions.

The subsequent error analysis used the same frozen predictions. We calculated Macro-F1 for individual contexts, tissues, studies and pair target-coverage groups, always using the same three-label definition. Macro-F1 was calculated over the same three predefined classes for every subgroup; classes absent from a subgroup therefore contributed an F1 of zero. For candidate prioritization, we ranked test observations by predicted synergy probability and calculated precision among the top 1%, 5% and 10%, using ceiling rounding for list size and sample-identifier order to resolve ties. Enrichment was precision divided by test synergy prevalence. The rankings concern observed pair–context–study records. They were not used to select checkpoints or tune thresholds.

Calibration was examined using ten equal-width probability bins, multiclass Brier score and log loss. As a descriptive probability reference, a constant predictor assigned the training class frequencies to every test observation. No recalibration was fitted. Because classification, ranking and calibration measure different properties, representation rankings were reported separately for each objective.

### Additional program and biological analyses

A separate completed experiment transformed Hallmark scores into NMF programs with K in {5, 10, 15, 20, 30}. Min–max scaling, NMF fitting and subsequent standardization used training contexts; held-out values were clipped to the training min–max range. Four combinations of programs with E and R were trained in three settings with five seeds, giving 300 candidate trainings. K was selected by validation Macro-F1v within each condition and seed, leaving 60 selected test evaluations. We analyzed their retained summaries.

A descriptive analysis examined regulon patterns associated with an expression-defined senescence-like axis in DepMap. Its score was the mean standardized expression of CDKN2A, CDKN1A and SERPINE1 minus that of MKI67, PCNA, TOP2A, MCM2, MCM5 and CCNB1. The top and bottom 20 profiles were compared. This analysis used a separate contextualized CollecTRI network: prior signs were multiplied by absolute across-DepMap Spearman correlations, with a 0.1 magnitude threshold, yielding 580 transcription factors and 20,058 edges. Group comparisons used Mann–Whitney tests with Benjamini–Hochberg correction. This diagnostic was not used to train or select the combination predictor.

## Results

### A common evaluation dataset exposes distinct generalization tasks

The eligibility filter retained 47.24% of the 639,410 aggregated observations. No interaction accounted for 80.19% of the eligible dataset, compared with 8.48% antagonism and 11.33% synergy (Figure 2). This imbalance makes overall accuracy an incomplete description of performance. The test synergy prevalence also differed across settings: 11.66% for unseen lines, 10.54% for unseen drugs and 14.17% for the joint holdout.

**Figure 2.**
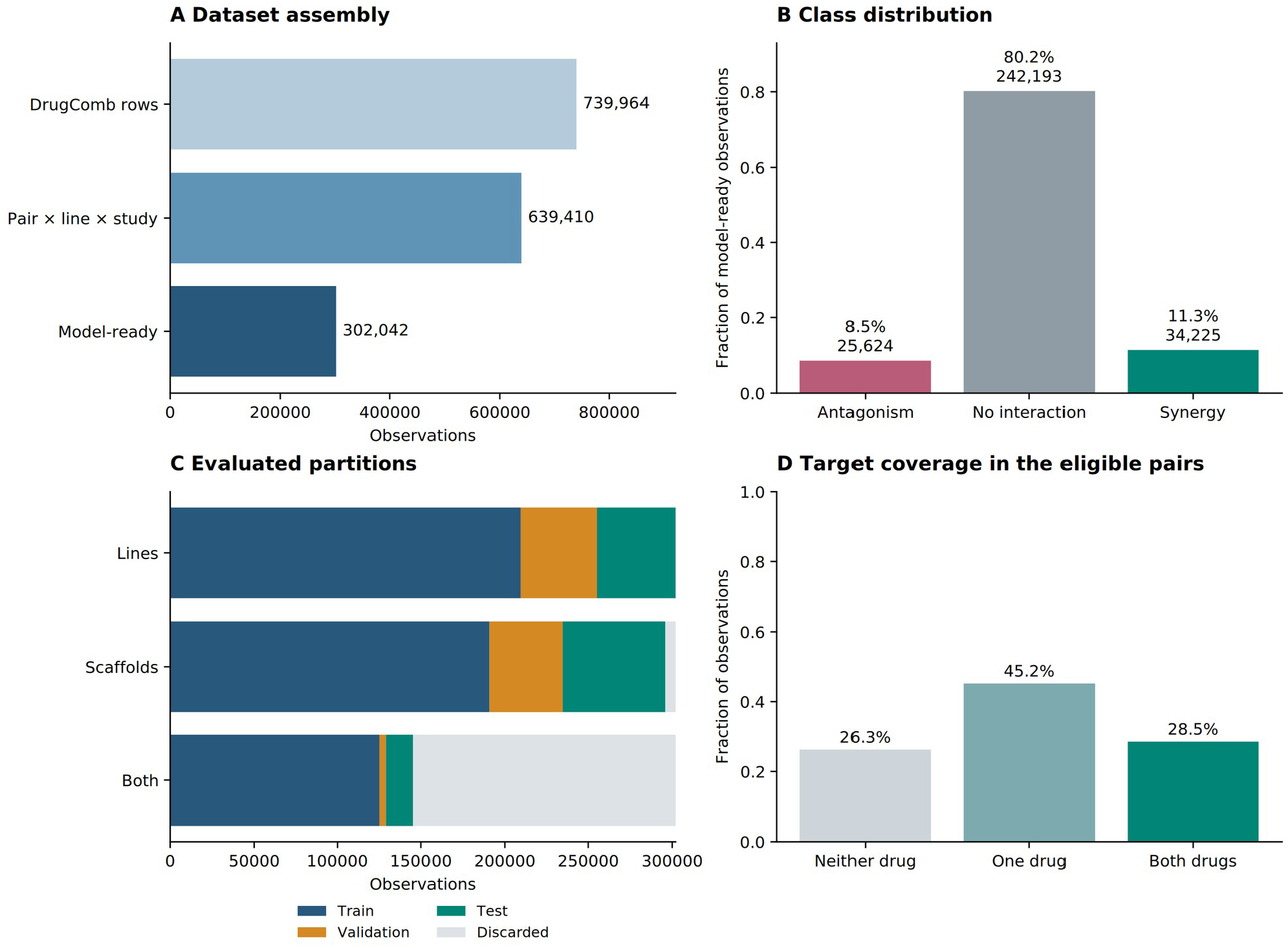
Dataset composition and evaluation partitions. A, progression from raw DrugComb rows to aggregated and model-ready observations. B, class proportions and counts in the common eligible dataset. C, training, validation, test and discarded observations under the three split rules. D, the fraction of eligible observations with zero, one or two drugs having at least one target mapped to HuRI. Drug-group holdouts require at least one held-out partner; the full rules and counts appear in Table 1.

**Table 1.**
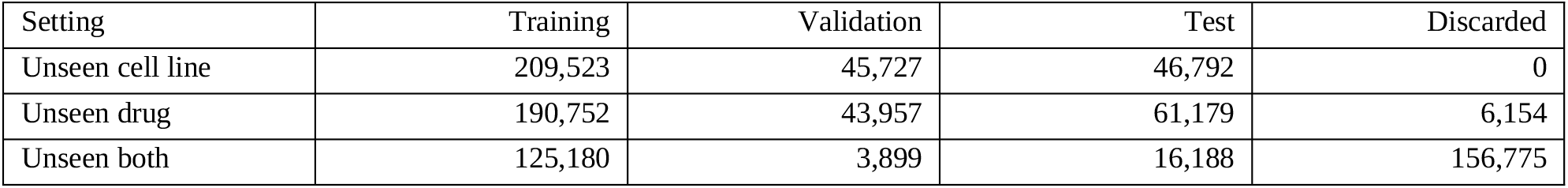
Observation counts by partition. The cell-line and unseen-both settings each use 109 training, 23 validation and 23 test contexts. The unseen-drug setting permits cellular contexts to recur across partitions. Observations whose group memberships did not satisfy the defined training, validation or test criteria were excluded from the corresponding analysis.

| Setting | Training | Validation | Test | Discarded |
| --- | --- | --- | --- | --- |
| Unseen cell line | 209,523 | 45,727 | 46,792 | 0 |
| Unseen drug | 190,752 | 43,957 | 61,179 | 6,154 |
| Unseen both | 125,180 | 3,899 | 16,188 | 156,775 |

Target coverage varied at the level of tested pairs. Neither drug had a mapped HuRI target in 26.3% of eligible observations, one drug had a mapped target in 45.2%, and both did in 28.5%. The target-availability indicators explicitly distinguished these coverage states, allowing drugs without HuRI-mapped targets to be represented without excluding the corresponding observations. Cellular context representations were complete by construction, and the same eligibility filter was used for all seven representations, so comparisons among E, P and R combinations were not confounded by differences in eligible observations.

The unseen-both holdout retained fewer training observations and excluded 156,775 records that crossed the designated context and drug partitions. It therefore probes a different training and test distribution from either single-axis holdout. Absolute differences between settings combine the effect of the held-out information with changes in sample size, class composition and the combinations retained by the split rules and should not be interpreted as a controlled comparison of task difficulty.

### The highest-scoring representation depends on the held-out information

P+R achieved the highest mean Macro-F1 for unseen cell lines (0.4737 ± 0.0077) compared with 0.4503 ± 0.0152 for E (Table 2; Figure 3). For unseen drugs, E+R achieved the highest mean (0.5583 ± 0.0101),followed closely by P+R (0.5565 ± 0.0043). In the unseen-both setting, P alone achieved the highest mean Macro-F1 (0.4225 ± 0.0233). Thus, no single representation performed best across all three generalization settings, and the full E+P+R representation did not lead in any setting.

**Table 2.** Test Macro-F1 across cellular representations. Values are means ± sample SD over five training seeds on the same partition within each setting. Bold identifies the largest observed mean in each column, rather than a statistically established winner. E, expression; P, pathways; R, regulons.

| Representation | Unseen cell line | Unseen drug | Unseen both |
| --- | --- | --- | --- |
| E | $0.4503 \pm 0.0152$ | $0.5494 \pm 0.0092$ | $0.4032 \pm 0.0275$ |
| P | $0.4584 \pm 0.0081$ | $0.5336 \pm 0.0177$ | **$0.4225 \pm 0.0233$** |
| R | $0.4526 \pm 0.0236$ | $0.5479 \pm 0.0172$ | $0.4107 \pm 0.0146$ |
| E+P | $0.4529 \pm 0.0159$ | $0.5505 \pm 0.0089$ | $0.3976 \pm 0.0155$ |
| E+R | $0.4535 \pm 0.0281$ | **$0.5583 \pm 0.0101$** | $0.4073 \pm 0.0164$ |
| P+R | **$0.4737 \pm 0.0077$** | $0.5565 \pm 0.0043$ | $0.4009 \pm 0.0058$ |
| E+P+R | $0.4474 \pm 0.0179$ | $0.5471 \pm 0.0126$ | $0.3992 \pm 0.0145$ |

**Figure 3.**
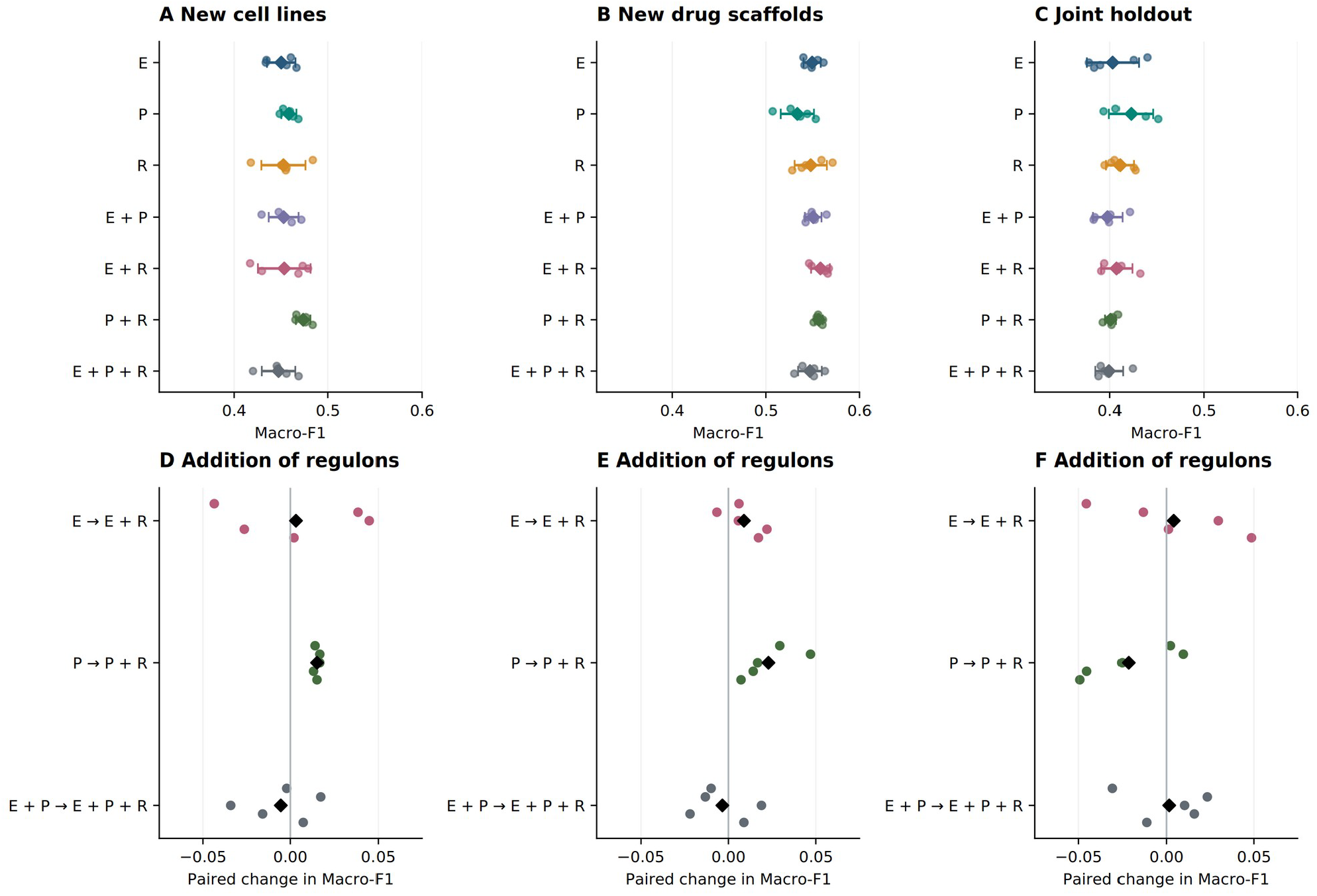
Representation performance and paired regulon additions. A–C, test Macro-F1 for all seven representations. Small circles indicate the five runs; diamonds and horizontal bars indicate the mean and sample SD. D–F, matched-seed changes when R is added to E, P or E+P; black diamonds indicate the mean difference. Each column uses one fixed partition. Positive changes indicate higher Macro-F1 after adding R. E, expression; P, pathways; R, regulons.

Performance was not explained simply by representation dimensionality or total model size. P contains 50 input features, compared with 771 for R and 942 for E. P+R contains 821 input features and slightly fewer trainable parameters than the E model, yet achieved higher mean Macro-F1 for unseen cell lines.

Conversely, adding R to a fixed base representation necessarily introduces an additional encoder branch and increases the number of trainable parameters relative to that base model. The present design therefore does not isolate the contribution of biological information from that of increased model capacity. These comparisons quantify the performance of the implemented representations but do not identify a unique causal explanation for those differences.

### Regulons show conditional complementarity with pathways

Across training seeds, the most consistent positive addition was P→P+R. Adding R to P increased Macro-F1 by 0.01527 ± 0.00159 for unseen cell lines and by 0.02289 ± 0.01566 for unseen drugs, with positive paired differences for all five paired seeds in both settings. In the unseen-both setting, the mean difference reversed to −0.02159 ± 0.02685. Adding R to E produced smaller mean changes of +0.00322, +0.00888 and+0.00413 across the three settings, respectively, whereas adding R to E+P changed the means by −0.00545, −0.00346 and +0.00155.

For both P→P+R comparisons with positive differences across all five training seeds, the exact two-sided Wilcoxon p-value was 0.0625, with a Holm-adjusted value of 0.5625 across the nine prespecified contrasts. No Holm-adjusted comparison reached p < 0.05. Because the five runs within each setting used the same held-out partition, these comparisons quantify the magnitude and consistency of changes arising across training seeds rather than variation across independently sampled test sets. Inference about variation across held-out datasets would require repeated independent held-out assignments.

Effects also varied among the 23 held-out cell lines. The per-line Macro-F1 difference between P+R and P was positive for 17 of 23 lines, with median +0.00899 and a range from −0.07083 to +0.06187 (Figure 4C). The unweighted mean of the per-line differences was +0.00514, compared with +0.01527 for Macro-F1 calculated from all pooled test observations. These quantities weight the data differently and need not coincide: individual lines contributed between 43 and 9,452 observations, and seven lacked one of the three interaction classes. Thus, the improvement observed in the pooled test set was distributed across a majority of held-out lines but remained heterogeneous in magnitude and direction.

**Figure 4.**
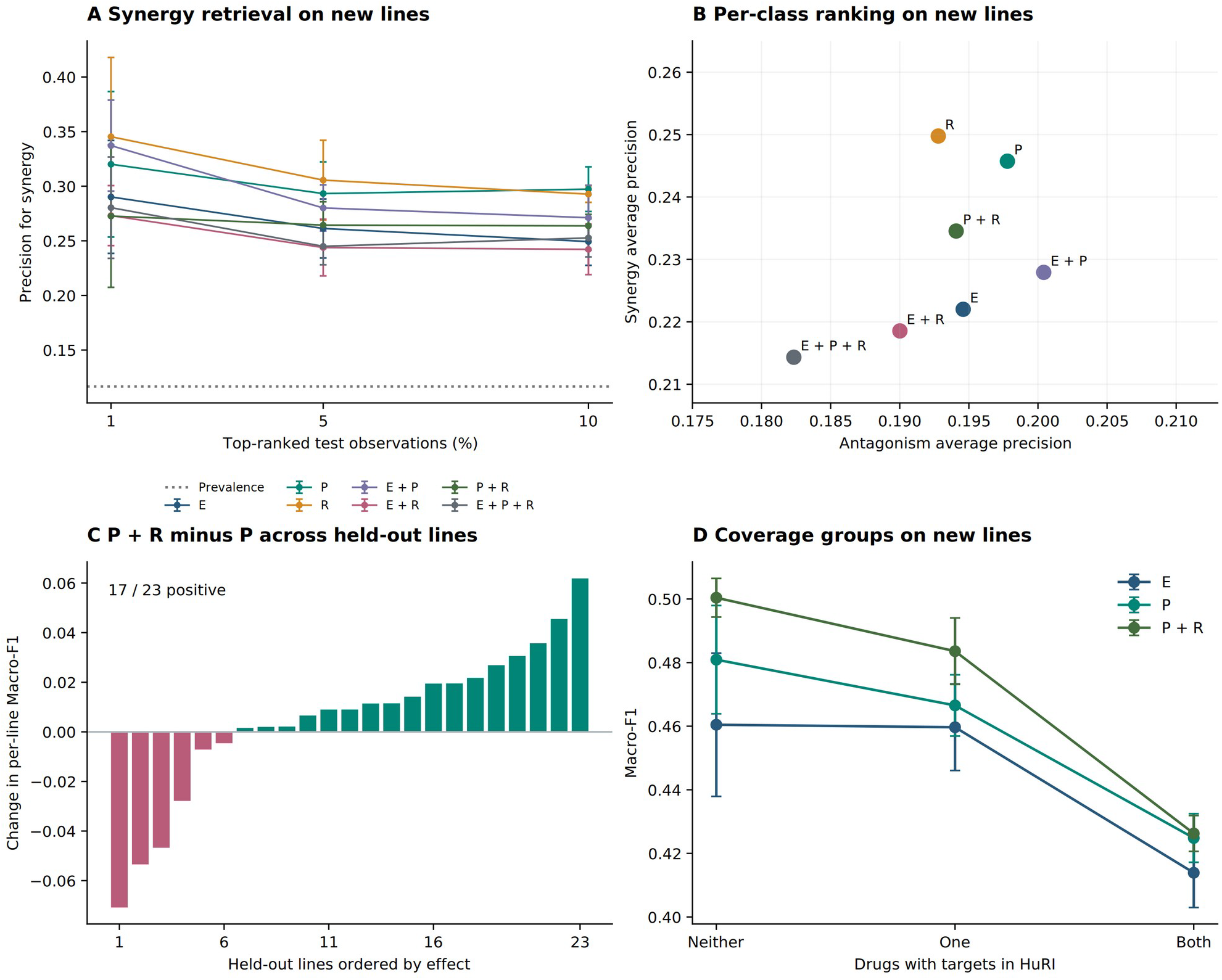
Error structure and candidate prioritization on unseen cell lines. A, mean synergy precision among the top 1%, 5% and 10% of test records ranked by predicted synergy probability; bars show training-seed SD and the dotted line shows class prevalence. B, mean AP for antagonism and synergy. C, P+R minus P differences in per-line Macro-F1, averaged over training seeds and sorted for display; seven of the 23 lines have two observed classes, and all use the fixed three-label definition. D, mean Macro-F1 ± SD by the number of drugs with mapped HuRI targets. The displayed E, P and P+R curves compare the expression reference with the direct pathway–regulon contrast; all seven variants are retained in the source tables.

### Classification and candidate retrieval favor different representations

The representation with the highest Macro-F1 for unseen cell lines did not maximize synergy ranking performance. R alone achieved synergy AP of 0.2498, compared with 0.2457 for P, 0.2345 for P+R and 0.2220 for E. Antagonism AP was highest for E+P at 0.2004, whereas P+R achieved 0.1941. Thus, improved performance under the three-class argmax decision rule did not necessarily correspond to improved ranking of test observations by predicted synergy probability.

The same distinction was evident in top-ranked retrieval (Figure 4A). Among the top 5% of unseen-cell-line test observations ranked by predicted synergy probability, mean synergy precision was 0.3056 for R, 0.2932 for P and 0.2644 for P+R. Relative to the test-set synergy prevalence of 0.1166, these values corresponded to 2.62-, 2.51- and 2.27-fold enrichment, respectively. Thus, the P+R gain in Macro-F1 did not translate into a higher synergy hit fraction at this screening budget. In the unseen-drug setting, P+R achieved top-5% precision of 0.5789, or 5.49-fold enrichment over the test prevalence of 0.1054, while E+R achieved 0.5551 despite having the highest mean Macro-F1. Full results for the three retrieval depths and all representations are provided with the numerical data.

Predicted class probabilities showed a different pattern. For unseen cell lines, mean multiclass Brier scores were 0.5742 for P+R and 0.5675 for P, compared with 0.3680 for a constant reference predictor that assigned the training-set class frequencies to every test observation. The training-frequency reference achieved lower Brier error than the evaluated models in all three generalization settings. This result does not contradict the observed ranking enrichment because the constant reference contains no within-class ranking information. Rather, it demonstrates that useful candidate ordering did not imply accurate probability estimates under the class-weighted training objective used here. Reliability plots are provided in Supplementary Figure S3.

Target-coverage strata further illustrated heterogeneity within the test distribution (Figure 4D). P and P+R achieved higher Macro-F1 among observations in which zero or one drug had a HuRI-mapped target than in the group in which both drugs had mapped targets. Because these strata contain different sets of compounds and observations, this comparison does not imply that target information reduces predictive performance. Instead, it indicates that nominal target-resource coverage alone is not a proxy for subgroup-level predictive performance. Tissue- and study-stratified results are included in the source tables.

### Additional pathway compression did not improve the best observed Macro-F1

The NMF analysis comprised 300 candidate training runs and 60 validation-selected test evaluations. For unseen cell lines, the highest mean Macro-F1 among the program-containing representations was 0.4661 for R+programs, compared with 0.4737 for P+R in the primary representation comparison. For unseen drugs, E+R+programs reached 0.5558, compared with 0.5583 for E+R. For the unseen-both setting, programs alone reached 0.4164, compared with 0.4225 for P. Thus, none of the validation-selected program representations exceeded the highest mean Macro-F1 observed in the corresponding setting of the primary comparison. These comparisons are descriptive and refer to the retained fixed partitions; complete seed-level results and the validation-selected values of K are provided in the Supplementary Data.

These results place a useful boundary on the value of additional functional compression. Hallmark pathway activities already provide a compact representation of expression-derived biological state, and further compression of these activities into NMF programs did not improve the highest observed Macro-F1 in any of the three generalization settings. This does not imply that lower-dimensional programs are intrinsically less informative: their performance depended on the representation with which they were combined and on the generalization setting. Rather, the results indicate that increasing the level of abstraction does not by itself guarantee improved predictive generalization.

### A separate expression-defined diagnostic links regulons to a senescence-like axis

We next asked whether the contextualized regulon activities captured regulatory patterns associated with an independently defined senescence-like expression axis in DepMap. The 20 DepMap profiles with the highest senescence-like expression scores and the 20 with the lowest scores were compared. The expression score was defined independently of the tested regulon activities from standardized expression of senescence/growth-arrest-associated and proliferation-associated markers, as described in Methods.

CEBPB and GATA4 regulon activities were higher in the high-score group, whereas E2F activity, defined as the mean standardized activity of E2F1–E2F8, was lower. The composite regulon score, defined as the mean of NF-κB, CEBPB and GATA4 activities minus E2F activity, was also higher in the high-score group. All four differences had Benjamini–Hochberg-adjusted p-values below 1.6 × 10^−7^. In contrast, NF4-κB activity, defined from RELA, RELB, REL, NFKB1 and NFKB2, showed no detected difference between groups (adjusted p = 0.4094). The complete distributions and tissue identities are provided in Supplementary Figure S5 and its source table.

These associations characterize an expression-defined axis within the available cancer cell-line panel rather than independently validating a senescent cellular state. Both group selection and regulon-activity inference depend on transcriptomic information, the high- and low-score groups have different tissue composition, and this diagnostic uses a contextualized regulatory network distinct from the fixed CollecTRI prior used in the drug-combination predictor. The analysis therefore provides biological context and a hypothesis-generating link to future senescence-focused applications, but does not demonstrate transfer to senescent primary cells or establish senolytic or senolytic-combination prediction performance.

## Discussion

This study identifies a conditional benefit of regulon-informed cellular representations for drug interaction prediction. The clearest matched comparison is the addition of R to P: the same architectural addition has positive mean effects on unseen lines and unseen scaffolds, and a negative mean effect when both axes are held out. Its impact is also heterogeneous among held-out lines and differs between Macro-F1 and synergy retrieval. These observations support selecting a representation in relation to the intended task rather than assuming that the largest set of biological features is the most transferable.

One interpretation is that pathways and regulons supply complementary summaries of basal expression. Hallmarks group genes by coherent processes, whereas signed regulons organize them by transcriptional control. Their combination may retain useful distinctions that either summary loses. However, the present experiments do not isolate that mechanism. Adding a branch also changes optimization, regularization and capacity, and the training data represent unevenly sampled biological and chemical spaces. The limited benefit of adding R to E or E+P further argues against treating regulon information as universally additive. Future matched-capacity and prior-perturbation controls can separate these explanations.

The comparison with prior work clarifies the scope of the contribution. Functional cell representations, including TF-based descriptions, are already part of the drug-synergy literature (Hosseini and Zhou 2023). Here, continuous fixed-prior scores and their complete combination with expression and pathways are examined inside one prediction framework. The useful finding is the pattern of complementarity and tradeoffs under different holdouts. Our data do not support a state-of-the-art claim against external models, because those models were not trained and evaluated on the same processed observations and assignments. The current result is a controlled representation study that can inform such a benchmark.

The separation between classification and retrieval has direct implications for experimental prioritization. Macro-F1 weights the three classification outcomes equally, while a screening campaign may primarily value the fraction of synergistic observations among a short candidate list. Those objectives can prefer different representations even when they use the same predictions. The R-only model’s stronger unseen-line synergy retrieval, together with the P+R model’s stronger Macro-F1, provides an example. A prospective application should therefore choose its model and decision criterion on validation data according to the intended experimental budget and the costs of false positives and false negatives.

Calibration adds another requirement. Class weighting changes the loss contribution of common and rare labels, and raw softmax outputs need not represent the prevalence-calibrated probability of observing a label in a new screen. Our Brier and reliability analyses make this issue visible. A downstream system could use rankings while separately developing probability calibration in an appropriate validation population. It should also consider the magnitude of the biological response: an interaction category alone cannot specify efficacy, toxicity or clinical benefit. More generally, benefits of combination therapy can arise through mechanisms beyond pharmacological synergy (Palmer and Sorger 2017).

For aging research, the main opportunity is to describe a cellular state using a stable set of interpretable biological coordinates and then test whether those coordinates support response prediction outside the oncology training domain. Prior work shows that TF and pathway activity estimation can be investigated across bulk and single-cell settings (Holland et al. 2020). This does not make a basal cancer-line score interchangeable with a donor-specific senescent profile. Differences in measurement platforms, gene coverage, cell composition and the biology of senescence can affect both the representation and its relationship to response. Transfer to senescence-focused applications therefore remains a hypothesis requiring direct evaluation in appropriate human cellular states and matched controls.

A focused next experiment would compare senescent and matched control cells of the same origin, measure combination dose responses, and distinguish interaction from selective activity. Senolytic selectivity is particularly useful because it retains a connection to viability-based training while introducing a clinically relevant contrast between cellular states. Senomorphic effects, including changes in secretory activity or cellular function, would require different response labels. The existing expression-defined diagnostic provides a way to inspect selected programs, but independent state annotation and response measurement remain necessary for this next step.

Several aspects of the evaluation determine how broadly the present results can be generalized. There is one fixed assignment per holdout setting, and the five seeds characterize variation in training. The split procedure uses class proportions when choosing assignments. Study-specific observations can share the same pair–context features within a partition, and the test populations differ between settings. In addition, the comparison uses a shared multi-modal eligibility filter and does not include trained drug-only, tissue-only, one-hot or matched-capacity controls. These choices are reported so that the next experimental stage can target the sources of uncertainty most directly: repeated group assignments, simple baselines, study-aware evaluation and independently annotated cellular states.

The NMF analysis further constrains the interpretation of abstraction. It provides no evidence that progressively higher-level compression alone yields progressively better generalization. Developing additional representations should therefore be tied to a specific biological or statistical hypothesis. For example, perturbation-response supervision could test whether observed drug-induced changes add information beyond basal state, while independently annotated senescent/control profiles could test whether the chosen regulons resolve the state relevant to selective response. These questions are distinct from increasing model depth or maximizing an existing test score.

In conclusion, fixed-prior regulon activities improve predictive performance in specific combinations of cellular representations and under specific forms of domain shift, but their value is neither universal nor independent of the evaluation objective. Pathway and regulon activities were complementary for some generalization tasks, whereas additional features or further pathway compression did not consistently improve performance. Classification, candidate retrieval and probability estimation also favored different representations, emphasizing that cellular-context encoding should be evaluated in relation to its intended use. By defining both the observed gains and their boundaries, this study provides a controlled empirical basis for selecting functional cellular representations and a concrete framework for subsequent response-based validation in aging-associated cellular states.

## Data and code availability

The accompanying reproducibility package contains the model source snapshot, exact retained feature definitions, split metadata, run metrics, publication analysis scripts and numerical sources for all figures. It also identifies the audited input files by SHA-256 and documents how the publication outputs are regenerated from saved predictions. DrugComb, DepMap, HuRI, Hallmark and CollecTRI are cited at their primary sources. Large input tables, per-observation prediction files and model checkpoints are distinguished from the compact publication package in its manifest.

