## Supplementary material: methods, tables and figures for "Regulon-informed cellular representations reveal task-dependent generalization in drug combination prediction"

### Supplementary Methods

#### S1 Feature construction and model dimensions

The feature definitions are fixed across the seven context conditions. Drug-target availability is an input feature, so lack of a mapped target is not treated as lack of a matrix row. By contrast, the common observation filter requires every context representation, even when a particular model uses only one. This preserves the sample population when comparing representations.

| Component | Input | Hidden and output dimensions |
|---|---:|---|
| Chemical branch | 2,048 | 512 → 128 |
| Direct-target branch | 8,245 | 256 → 32 |
| HuRI propagation branch | 8,245 | 512 → 128 |
| Availability | 3 | Identity |
| Drug fusion | 291 | 256 → 128 |
| Expression branch | 942 | 256 → 64 |
| Pathway branch | 50 | 64 → 64 |
| Regulon branch | 771 | 256 → 64 |
| Context fusion | 64 × active branches | 256 → 128 |
| Symmetric pair classifier | 512 | 256 → 128 → 3 |

**Supplementary Table S1. Dimensions of the shared model components.** Drug A and drug B use shared encoder parameters. Context conditioning combines projections, feature-wise modulation, an explicit drug–context interaction network and a gated residual. The classifier receives the sum, absolute difference and elementwise product of the conditioned drug vectors, together with the context vector. Dropout is 0.20. The full layer sequence is retained in the source snapshot.

| Context | Trainable parameters |
|---|---:|
| E | 8,498,467 |
| P | 8,248,035 |
| R | 8,454,691 |
| E+P | 8,522,403 |
| E+R | 8,729,059 |
| P+R | 8,478,627 |
| E+P+R | 8,752,995 |

**Supplementary Table S2. Model sizes.** Parameter counts come from the saved run summaries. Branch additions change capacity; the comparison does not include parameter-matched randomized-feature controls.

Hallmark and CollecTRI scoring use their corresponding measured gene universes rather than only the 942 expression-branch landmarks. In the documented ULM formulation, expression y for one sample is regressed across genes against the set weights x, with an intercept: $y = \beta_0 + \beta_1 x + \varepsilon$. The reported score is the t-statistic of $\beta_1$. Hallmark membership supplies unsigned weights, whereas CollecTRI supplies signed regulatory weights. This is a statistical activity estimate derived from the transcriptome; it is not a direct assay of a transcription factor's biochemical activity.

The fixed CollecTRI snapshot contains 41,714 retained interactions and 771 scored factors. The separately contextualized diagnostic network contains 20,058 interactions and 580 factors. A separate contextualized CollecTRI network was used only for the senescence-like descriptive analysis. For this network, CollecTRI regulatory weights were reweighted by the absolute Spearman correlation between TF and target expression across the DepMap panel, while retaining the direction of the prior regulatory interaction. Self-interactions were removed, interactions with correlation magnitude below 0.1 were excluded, and no interactions absent from the CollecTRI prior were introduced. The resulting contextualized network contained 20,058 interactions and 580 transcription factors.

For the senescence-like diagnostic, contextualized ULM activities for selected transcription factors were standardized across all shared DepMap profiles. NF-κB activity was defined as the mean standardized activity of RELA, RELB, REL, NFKB1 and NFKB2, whereas E2F activity was defined as the mean standardized activity of E2F1–E2F8. CEBPB and GATA4 were represented by their respective standardized regulon activities. A composite regulon score was calculated as the mean of NF-κB, CEBPB and GATA4 activity minus E2F activity. These diagnostic quantities were not used as features in the drug-combination predictor.

### S2 Training and partition reconstruction

All cellular-context models were trained using AdamW with a learning rate of $10^{-4}$, weight decay of $10^{-4}$, batch size 128 and a maximum of 50 epochs. Weighted cross-entropy was used, with class weights calculated from the training partition as $N/(3n_c)$, where $N$ is the number of training observations and $n_c$ is the number belonging to class $c$. Validation Macro-F1 was used for checkpoint selection and early stopping with a patience of eight epochs. A ReduceLROnPlateau scheduler also monitored validation Macro-F1, with a reduction factor of 0.5 and patience of two epochs. Gradient norm was clipped at 5. Training seeds were 42–46. CUDA deterministic algorithms were not enforced, so the five runs characterize training stochasticity under the fixed data assignment rather than deterministic repetitions.

For the publication analysis, scaffold groups were independently reconstructed from the retained canonical molecular structures using RDKit. Cyclic molecules were grouped by Bemis–Murcko scaffold without chirality. Acyclic structures use the full canonical SMILES. When scaffold generation produced an empty scaffold, the full canonical structure was used as the grouping identifier. Across the 3,042 drugs in the common eligible population, this procedure yielded 1,942 scaffold groups, with no ambiguous multiple-structure drug identifiers. Partition membership lists were then read from the original split JSON files rather than regenerating or reselecting the partitions. Reconstructed observation counts and test observation identities matched those stored with the saved predictions.

For scaffold-based holdouts, a test observation was required to contain at least one drug assigned to a designated test scaffold group and no drug assigned to a validation scaffold group; the other drug could belong to a training scaffold group. Thus, the unseen-drug setting requires at least one held-out drug rather than requiring both members of a combination to be unseen. In the joint setting, a drug identity can also be absent from the realized training observations because its observed combinations do not occur within training cellular contexts, even when its scaffold was not assigned to a held-out drug group. The identity audit therefore tests canonical-structure overlap specifically between the designated held-out scaffold groups and training structures. No such overlap was detected.

The split JSON files preserve the chosen group assignments, requested and selected seeds, class-composition criteria and observation counts. The selected seeds were 79 for the unseen-cell-line setting, 47 for the unseen-drug setting and 60 for the joint unseen-both setting. Each generalization setting used the same train–validation–test assignment for all five training seeds. Consequently, variation across the five runs reflects training stochasticity on a fixed partition and does not estimate variability across independently sampled holdout assignments. Validation and test class supports are included in the numerical split-audit table.

### S3 Metric definitions and subsequent analyses

Macro-F1 is the arithmetic mean of the F1 scores for antagonism, no interaction and synergy. Labels are predicted by argmax of the stored class probabilities. Balanced accuracy averages class recalls. AP is computed separately for each class against all others; macro AP averages those three values. It uses the stepwise average-precision estimator rather than trapezoidal integration of a precision–recall curve. The original output column called AUPRC corresponds to this AP implementation.

The independent verification requires exact identifier and label agreement, finite probabilities in [0, 1], probability sums within $2 \times 10^{-6}$ of one, and agreement between stored predicted classes and argmax. Metrics are compared with the saved summaries using a $2 \times 10^{-5}$ tolerance to accommodate rounding of probabilities in the TSV exports. F1 and balanced accuracy match exactly. Scalar metrics use every observation. For display only, supplementary ROC/PR curves use up to 250 evenly spaced indices from the full curve arrays.

There are nine direct regulon-addition comparisons. For each, the five differences are paired by training seed. Exact two-sided Wilcoxon p-values are adjusted jointly by Holm's method. The two all-positive P → P+R comparisons have unadjusted $p = 0.0625$ and adjusted $p = 0.5625$. Other adjusted p-values equal 1. These are comparisons of training repetitions under fixed partitions. We do not bootstrap individual rows to represent uncertainty over new donors, cell lines or drug groups.

Per-context, tissue, study and target-coverage summaries use the same fixed three-class F1 definition and report class support. Top-fraction analysis ranks all test records within a run by predicted synergy probability, takes $\mathrm{ceil}(f \times N)$ records for $f = 0.01$, 0.05 or 0.10, and reports their observed synergy fraction. Sorting sample IDs before stable score ordering fixes tie handling. Study-specific repeats remain records, so the lists are not unique-drug-pair lists.

Multiclass Brier score is the mean sum over the three squared probability errors, with range [0, 2]. Log loss is the mean negative log probability of the observed class. The constant reference predicts training class frequencies, so its test Brier score is $1 + \Sigma q^2 - 2\Sigma pq$, where q denotes training frequencies and p denotes test frequencies. The resulting values are 0.3680, 0.3181 and 0.3969 for the line, drug and joint settings. This reference evaluates probability error and has no ranking ability. For classification, predicting the training-majority class for every observation gives test Macro-F1 values of 0.2922, 0.2994 and 0.2879 in the same three settings. These analytical no-fit references do not replace trained baseline models. Calibration plots use ten fixed bins per class. Pooled curves concatenate counts from the five prediction sets for descriptive display, retaining repeated predictions of the same observations; they are not an ensemble or five independent test samples.

### S4 NMF program experiment

The program experiment compressed the 50 Hallmark activity scores using non-negative matrix factorization (NMF) with K = 5, 10, 15, 20 or 30 NMF components. Before NMF, Hallmark activities were min–max scaled using parameters estimated from the training cellular contexts only, producing non-negative training values. Validation and test activities were transformed using the same training-derived parameters and clipped to the training range before projection. NMF was fitted exclusively on the training contexts. The resulting program features were subsequently standardized using parameters estimated from the training data and applied unchanged to validation and test data. The program experiment used the same train–validation–test assignments as the corresponding primary representation experiments.

Four program-containing cellular-context representations were evaluated: programs alone, E+programs, R+programs and E+R+programs. Five values of K, four representations, three holdouts and five seeds yield 300 candidate training runs. Within each representation, generalization setting and training seed, K was selected using validation Macro-F1 only; test performance was not used for component-number selection. This procedure produced 60 validation-selected models for test evaluation. Test metrics are summarized as means and sample standard deviations across the five training seeds. For context, the highest mean Macro-

F1 obtained among the seven representations in the primary experiment is shown for each generalization setting as a descriptive reference; no additional significance test was performed against this reference.

| Program representation | Unseen cell line | Unseen drug | Unseen both |
|---|---|---|---|
| Programs | 0.4629 | 0.5316 | 0.4164 |
| E+programs | 0.4642 | 0.5533 | 0.3930 |
| R+programs | 0.4661 | 0.5480 | 0.4098 |
| E+R+programs | 0.4614 | 0.5558 | 0.4024 |

**Supplementary Table S3. Validation-selected NMF-program test Macro-F1.** Means over five training seeds. The complete table includes SD, additional metrics and the selected K for each run.

### S5 Expression-defined senescence-like diagnostic

Default DepMap profiles shared between the expression and contextualized-regulon activity matrices were ranked using the expression-defined senescence-like score described in the main Methods. Briefly, expression of each marker was standardized across all shared profiles, and the score was calculated as the mean standardized expression of CDKN2A, CDKN1A and SERPINE1 minus the mean standardized expression of MKI67, PCNA, TOP2A, MCM2, MCM5 and CCNB1. The 20 profiles with the highest scores and the 20 with the lowest scores were designated senescence-like and non-senescent-like, respectively.

Contextualized ULM activities for the selected transcription factors were standardized across the same shared DepMap profiles. NF-κB activity was defined as the mean standardized activity of RELA, RELB, REL, NFKB1 and NFKB2, and E2F activity as the mean standardized activity of E2F1–E2F8. CEBPB and GATA4 were represented by their respective standardized regulon activities. A composite regulon score was defined as the mean of NF-κB, CEBPB and GATA4 activities minus E2F activity. The high- and low-score groups were compared for these five quantities using two-sided Mann–Whitney tests, with Benjamini–Hochberg correction across the five comparisons. Rank-biserial correlation was calculated as an effect-size measure, with positive values indicating higher activity in the senescence-like group. All 40 profile identities, expression-defined senescence-like scores, regulon-derived activities, composite regulon scores and group assignments are provided in the supplementary source data.

The high- and low-score groups differ in cellular origin and were not matched by lineage or tissue. In particular, the low-score group includes hematopoietic models, while the high-score group includes fibroblast-associated profiles. Consequently, differences between these groups may reflect a mixture of proliferation state, lineage and other transcriptomic differences rather than senescence alone. The analysis is therefore presented as a descriptive association with an expression-defined senescence-like axis rather than as independent validation of cellular senescence or as a drug-response benchmark. Group assignment and regulon-activity inference both depend on transcriptomic data, and the contextualized regulatory network was itself constructed using TF–target expression correlations across the DepMap panel. Neither the expression-defined group labels nor the contextualized regulon activities were used in training or selecting the drug-combination predictor.

### Supplementary figures

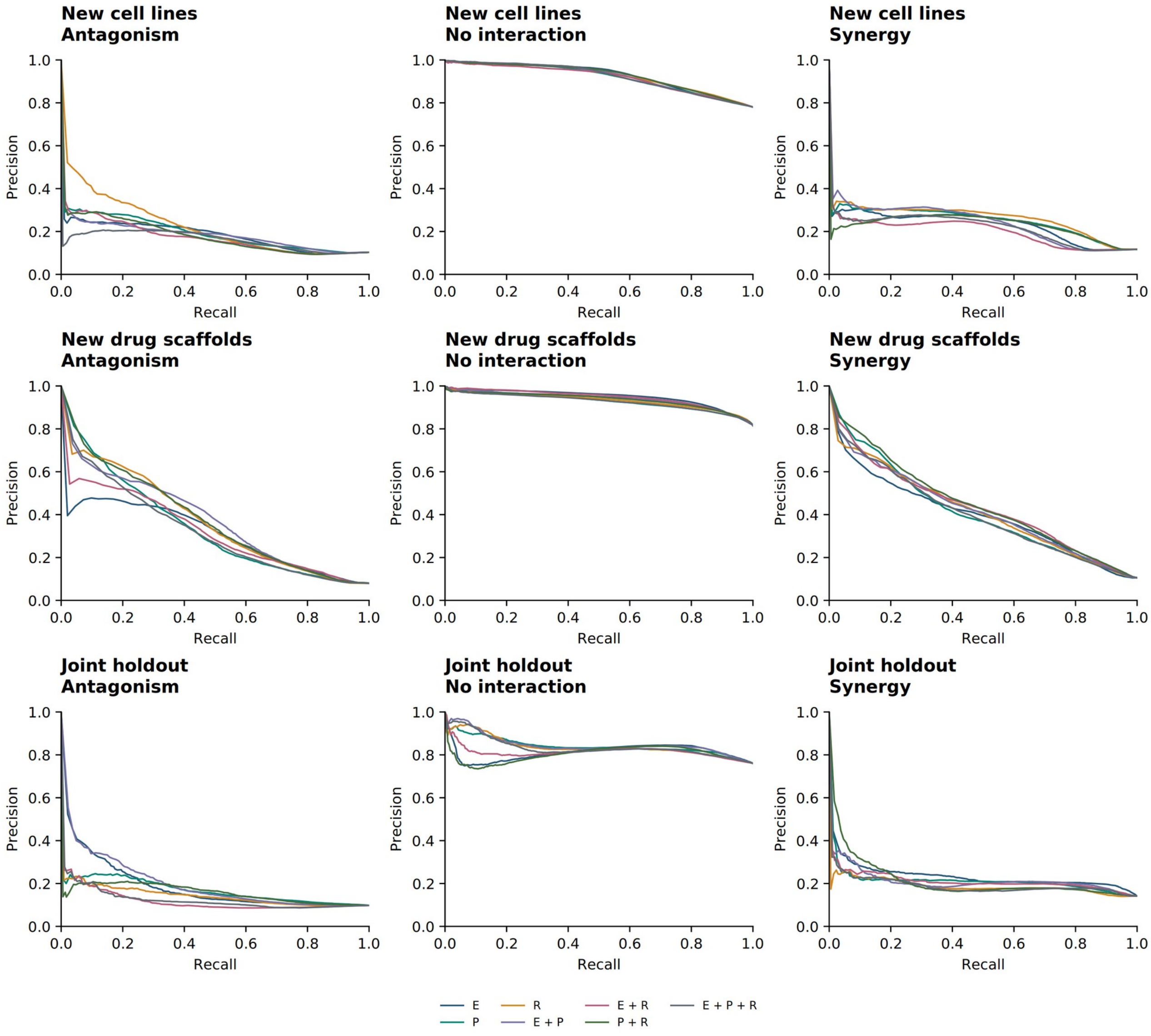


**Supplementary Figure S1. Precision–recall curves.** Curves are shown for each class and holdout setting for the predetermined seed 42. All seven context representations are included. Scalar AP results and all five seeds are provided in the source tables; the displayed seed is not selected by performance. E, expression; P, pathways; R, regulons.

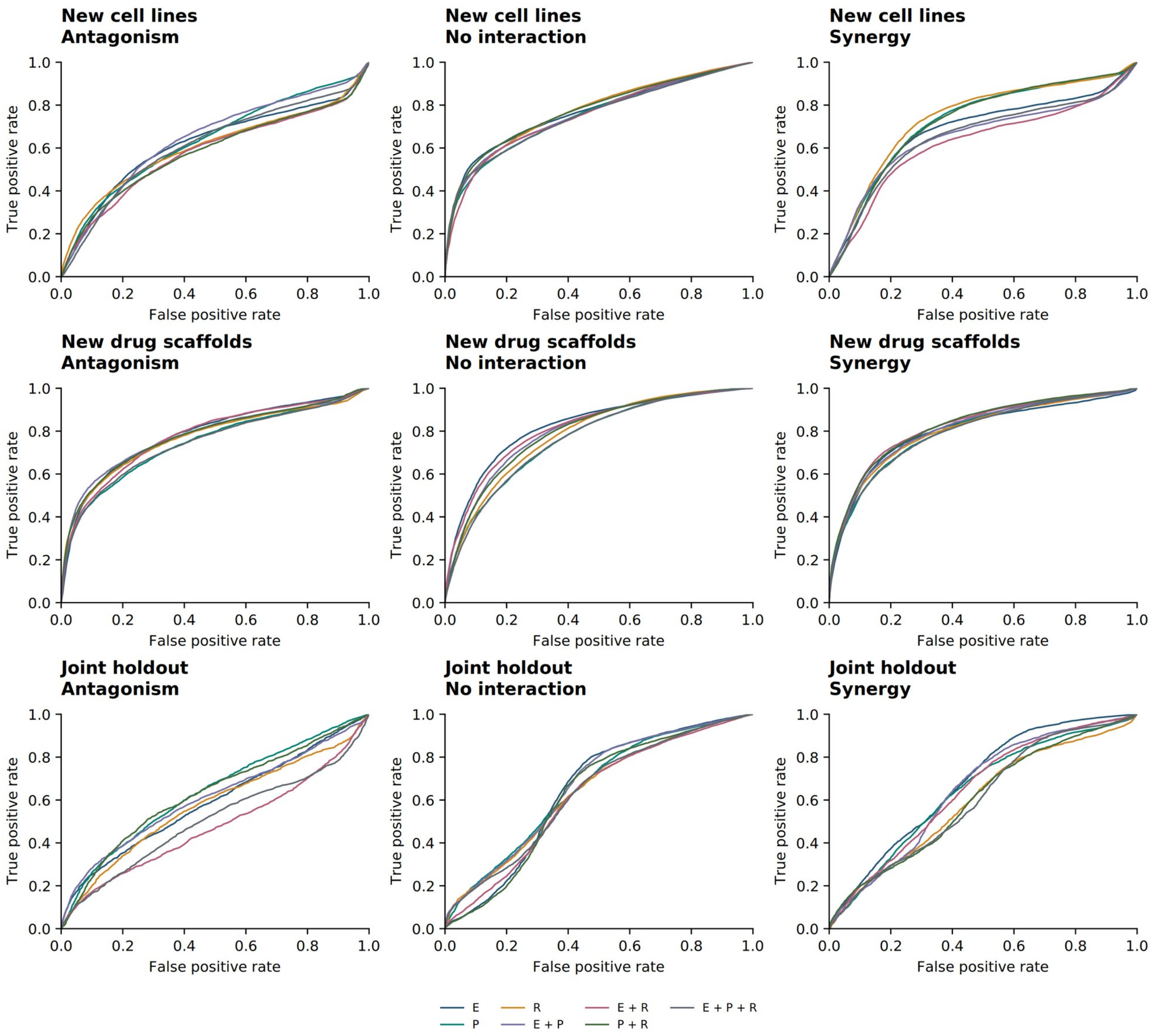


**Supplementary Figure S2. ROC curves.** One-versus-rest curves for each class and holdout setting, using seed 42 and all seven representations. The fixed-seed display complements the complete numerical results and does not estimate uncertainty across partitions.

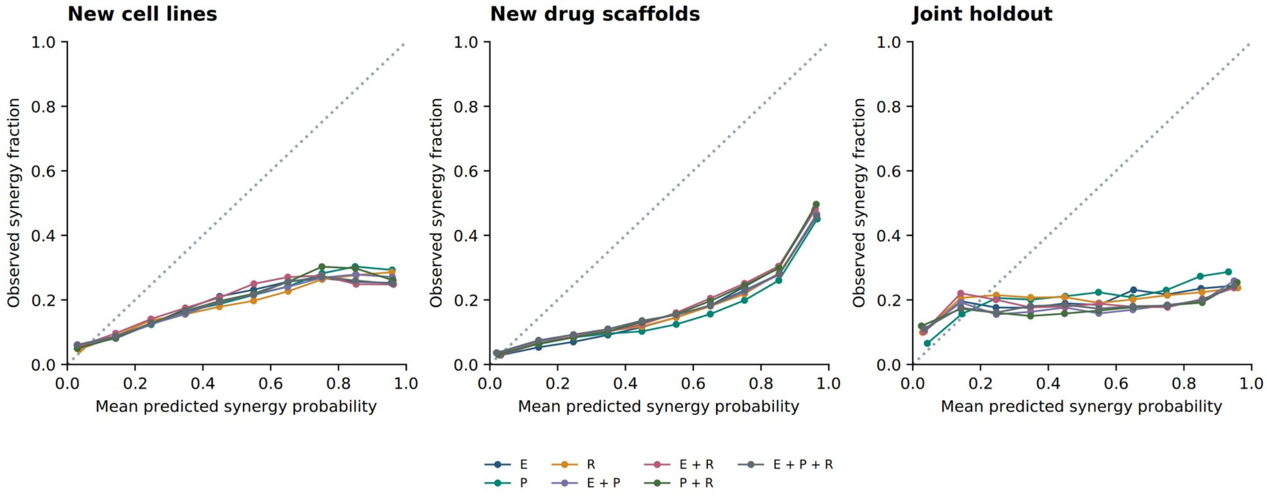


**Supplementary Figure S3. Reliability of predicted synergy probability.** Each point gives the mean predicted synergy probability and observed synergy fraction in a fixed probability bin, pooled descriptively across the five runs. The diagonal denotes calibration. Class weighting was used during training, and no probability recalibration was fitted. Full bin counts and per-run Brier/log-loss results are supplied.

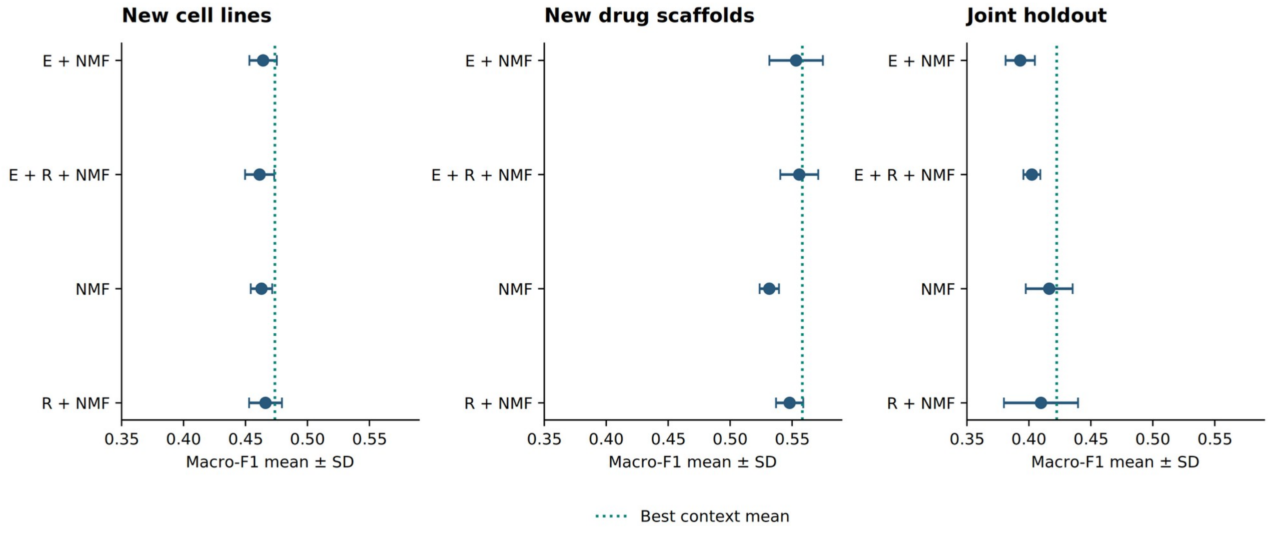


**Supplementary Figure S4. NMF program results.** Means ± sample SD of test Macro-F1 over five seeds after choosing K by validation Macro-F1. The vertical dotted line marks the largest mean among the seven original context representations for that holdout. E, expression; R, regulons; NMF, pathway-derived programs.

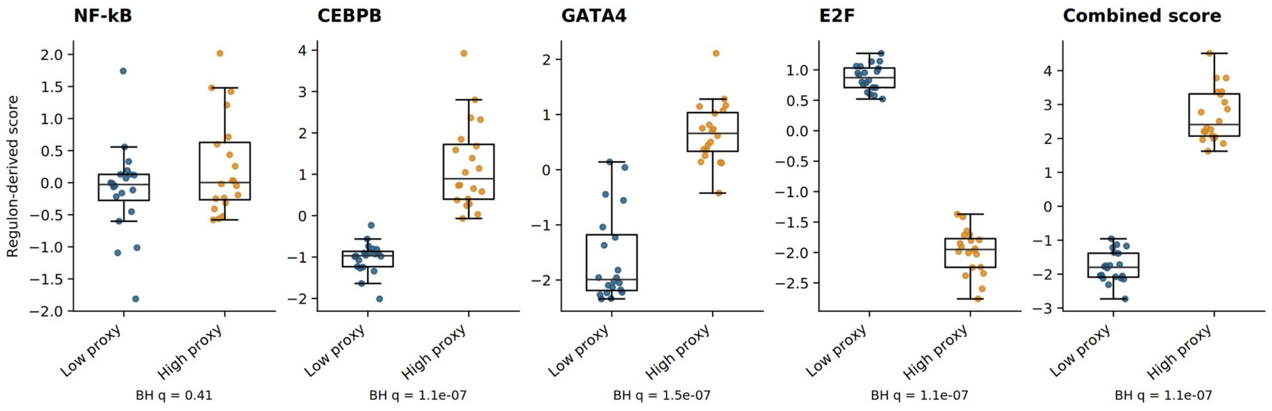


**Supplementary Figure S5. Regulon scores in expression-defined high- and low-proxy groups.** Each group contains 20 DepMap profiles. Points are individual profiles, boxes show the median and interquartile range, and whiskers extend to the most extreme values within 1.5 interquartile ranges; all observations remain visible as points. The two-sided Mann–Whitney comparisons use Benjamini–Hochberg adjustment across the five scores. NF-κB does not show a detected group difference. Group labels refer to proxy-defined profiles, rather than experimentally annotated senescent and matched control samples.

### Supplementary numerical data

The numerical package contains the following tables, with full precision retained in the machine-readable files:

- run_metrics.tsv and condition_summary.tsv: independently verified run-level metrics and aggregates.
- paired_regulon_effects.tsv and paired_regulon_statistics.tsv: matched-seed differences and the nine adjusted comparisons.
- split_audit.tsv, drug_groups.tsv and original split JSONs: group assignments, counts and identity checks.
- subgroup_metrics.tsv and strata_*.tsv: context, tissue, study and coverage results with sample/class counts.
- top_fraction.tsv and top_fraction_summary.tsv: precision and enrichment at each fraction for every run.
- reliability.tsv, probability_benchmarks.tsv and confusion.tsv: probabilistic diagnostics and error counts.
- curves_seed42.tsv and figure4_line_effects.tsv: numerical points used for the corresponding displays.
- Original context/program summaries, senescence-diagnostic tables, input manifests and upstream file hashes.

The model source snapshot preserves the implementation used for these experiments. The publication scripts reconstruct the analyses from saved observations and predictions without invoking the training or inference entrypoints. The exact original training-library environment was not recorded; the publication-analysis dependency versions and its verification results are supplied separately. Upstream resource identity is documented by retained feature definitions and input hashes where a named release is unavailable.
